# Trap-TRAP, a versatile tool for tissue-specific translatomics in zebrafish

**DOI:** 10.1101/2021.10.22.465445

**Authors:** Jorge Corbacho, Estefanía Sanabria-Reinoso, Ana Fernández-Miñan, Juan R. Martínez-Morales

## Abstract

Developmental and physiological processes depend on the transcriptional and translational activity of heterogeneous cell populations. A main challenge in gene expression studies is dealing with this intrinsic complexity while keeping sequencing efficiency. Translating ribosome affinity purification (TRAP) methods have allowed cell-specific recovery of polyribosome-associated RNAs by genetic tagging of ribosomes in selected cell populations. Here we combined the TRAP approach with adapted enhancer trap methods (trap-TRAP) to systematically generate zebrafish transgenic lines suitable for tissue-specific translatome interrogation. Through the random integration of the eGFP:rpl10a cassette, we have generated stable lines driving expression in a variety of tissues, including the retina, skeletal muscle, lateral line primordia, rhombomeres, or jaws. To increase the range of applications, a *UAS:TRAP* transgenic line compatible with available Gal4 lines was also generated and tested. The resulting collection of lines and applications constitutes a resource for the zebrafish community in developmental genetics, organ physiology and disease modelling.

## Introduction

The precise combination of repressed and activated genes determines the identity and transcriptional state of the cells, and thus controls their shape, mechanical properties, physiology, pathology, and survival. Genome-wide analysis of the transcriptome in different tissues provides very helpful information on the cells’ state through time. The emergence of Next Generation Sequencing (NGS) technologies allowed generating large volumes of sequencing data per run, offering whole-genome coverage at reduced costs. Among the numerous NGS applications, RNA-seq has become a method of choice in transcriptomics due to its high reproducibility, unbiased detection, single nucleotide resolution and quantitative estimation over a large dynamic range of gene expression ^1^. Despite its numerous advantages, RNA-seq analytical power is significantly diminished when complex tissues, such as the brain, are examined. Using an “en masse” approach, gene expression profiles cannot be assigned to any specific cell type, but reflect averaged gene expression across the entire tissue. Many of these disadvantages have been overcome by single-cell RNA sequencing (scRNA-seq) technologies, which permit the characterization of heterogeneous cell populations, making possible to detect the signature of rare cell types or transient cellular states ^2^. However, due to the low amount of starting material, scRNA-seq methods have their own technical limitations such as loss of spatial information, low capture efficiency, and frequent dropout events ^3^. In consequence, only a fraction of the transcriptome of each cell can be detected by scRNA-seq, and the technical noise is higher than in bulk RNA-seq ^4^. Therefore, the in-depth transcriptomic characterization of a given cell type still depends on our ability to isolate them either through micro-dissection or using fluorescence-activated cell sorting (FACS) protocols.

Unfortunately, severe dissociation procedures such as those required before FACS, may distort gene expression inducing a cellular response to stress ^5, 6^. As an alternative to flow cytometry, a number of approaches have been developed to directly label RNA, such as TU-tagging ^7^, or RNA-binding proteins ^8, 9^. Among them, the translating ribosome affinity purification (TRAP) technology stands out due to its low toxicity and its suitability in both, vertebrates and *Drosophila* ^8, 10, 11, 12^. This method is based on cell-type specific ribosome tagging, by expressing a GFP-tagged version of the large subunit ribosomal protein L10a (EGFP-Rpl10a) under the control of a tissue-specific promoter of choice. Then, labelled polyribosomes can be affinity purified to specifically pull-down associated mRNAs, which are a precise representation of translated genes in the cell population of interest. The TRAP methodology provides important advantages, as it does not require tissue fixation or dissociation and the cells of interest are marked, which facilitates in parallel imaging studies. Moreover, analysing the translating mRNA pool provides a closer representation of the protein content than the examination of the whole mRNA profile ^8, 13^.

In zebrafish, different laboratories have successfully implemented the TRAP approach using tissue-specific drivers. They include the promoters of the genes *tyrp1* for melanocytes ^14^, *cmlc2* for cardiomyocytes ^15^, *actc1b* for skeletal myocytes ^16^, or *lyz* for neutrophils ^17^. One of the main disadvantages of the TRAP methodology is precisely the need to create a customized transgenic line for each cell type to be analysed. This is a laborious and technically challenging procedure that may discourage many zebrafish laboratories; particularly when no suitable drivers are available for the population of interest.

To overcome some of these limitations, we have developed a strategy combining the TRAP technology with a traditional enhancer trap approach: here referred as *trap-TRAP*. Enhancer trap methodologies allow the random genomic insertion of a reporter gene to capture the activity of nearby cis-regulatory elements ^18^. In zebrafish, the adaptation of transposons, in particular that of the medaka element Tol2, has greatly improved the efficiency of enhancer trap approaches, allowing the generation of collections of stable enhancer trap lines ^19, 20, 21^. Using our *trap-TRAP* approach, which takes advantage of the Tol2 system to generate random insertions of a *eGFP*-*rpl10a* cassette, we have isolated 33 tissue specific lines in a pilot screen. Furthermore, by placing the eGFP-rpl10a fusion under the control of the UAS element, we have also combined the TRAP technique with the Gal4/UAS transcriptional activation system ^22^. This allows expanding the applicability of the TRAP method to the entire collection of Gal4 drivers available in zebrafish ^23, 24^. Taken together, our approaches allow the systematic generation of transgenic lines for efficient tissue-specific translatome interrogation in zebrafish.

## Results and Discussion

### Vectors to increase the scope of the TRAP technology in zebrafish

To expand the range of potential applications of TRAP methods in zebrafish we designed two different Tol2-based vectors ^20^. The first, *Tol2*_*trap:TRAP* (Figure 1A), allows combining translating ribosome affinity purification (TRAP) with a standard enhancer trap approach. To this end, the TRAP cassette (*eGFP*-*rpl10a*) was fused to the *gata2p* minimal promoter ^25^. The random integration of this vector in the zebrafish genome allowed capturing the activity of nearby cis-regulatory elements (i.e. enhancers), leading to the expression of *eGFP*-*rpl10a* in a tissue-specific manner. The second vector, *Tol2*_*UAS:TRAP* (Figure 1C), was designed to take advantage of the collection of Gal4 lines available in zebrafish ^26^ as drivers for the TRAP cassette. Both vectors were then tested in transgenic assays as described in the following sections.

**Figure 1.**
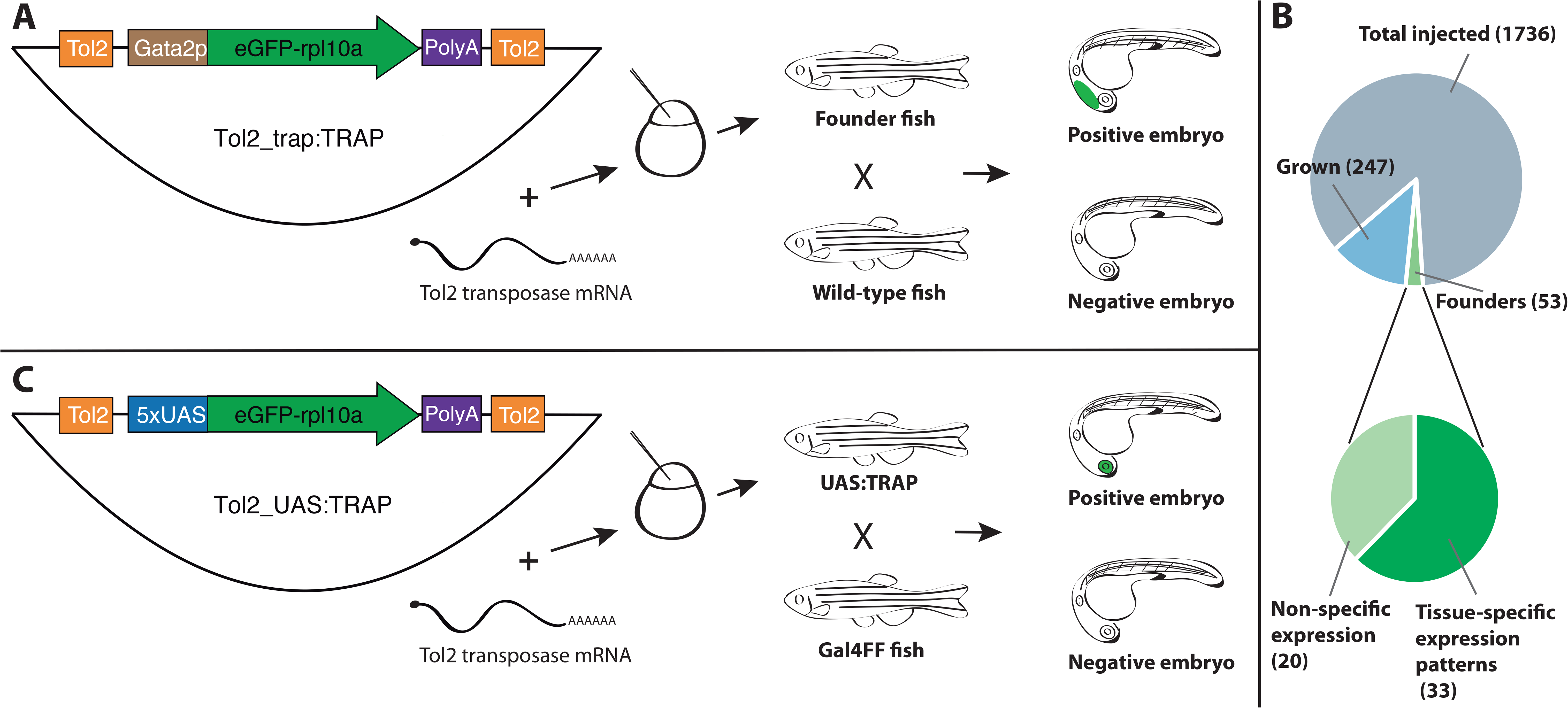
Diagrams of vectors and procedures used to expand TRAP approaches in zebrafish. A: The Tol2_trap:TRAP vector comprises a cassette containing the eGFP-rpl10a fusion gene (green) and the gata2p minimal promoter (brown). This cassette is flanked by Tol2 recognition sequences (orange). This vector was injected together with Tol2 transposase mRNA in one-cell stage zebrafish embryos. Once grown, adult fish were screened for eGFP-rpl10a expression in their progeny. B: Pie charts showing the efficiency rate of the trap:TRAP approach. C: The Tol2_UAS:TRAP vector comprises a cassette that contains the eGFP-rpl10a fusion gene (green) together with the 5xUAS element (blue). This cassette is also flanked by the Tol2 recognition sequences (orange). This vector together with *Tol2 transposase* mRNA were injected together in one-cell stage embryos. Once grown, adult fish were outcrossed with a Gal4 line to identify founders.

### Generation of the trap-TRAP transgenic lines

To test TRAP compatibility with an enhancer trap approach we carried out a pilot screen in zebrafish. The *Tol2*_*trap:TRAP* vector was injected in one-cell stage zebrafish embryos together with Tol2 transposase mRNA synthesized in vitro. Embryos showing any fluorescence at 24 hpf were selected and raised. A total of 300 adult fish were screened for eGFP-rpl10a expression by outcrossing them with wild-type animals. A minimum of 100 F1 embryos from each cross was examined at 24 and 48 hpf. We identified 53 positive founders, indicating a trapping efficiency of 17.5% (Figure 1B). This efficiency rate falls within the range of previously reported trapping screens using the Tol2 transposition system: 12% for enhancer trapping ^27^ and 23% for gene trapping ^28^. Among all founders, 33 (62% of total) showed tissue-specific expression patterns often restricted to a single domain. Each of these founders was isolated and outcrossed to generate stable lines, which were designated as TT followed by an identifier number. In most cases, the eGFP-rpl10a distribution indicated that the integration event was unique. However some founders showed segregation of expression patterns in their progeny, indicating that this event occurred more than once. This was the case for the lines TT1, which also included TT56; TT37, also including TT42, TT57, TT58 and TT59; TT48 also comprising TT50; and for TT53 that had a common founder with TT60. Among the stable lines, eGFP-rpl10a signal was detectable in a variety of tissues from different embryonic origin (i.e. from different germ layers) and located in different positions along the embryo axes. This observation indicates that the eGFP-rpl10a fusion does not compromise the random integration of the trapping cassette.

The expression patterns of the 10 most relevant transgenic lines (i.e. those exhibiting higher tissue specificity at 40-48 hpf) are shown in Figure 2. These include specific lines for hindbrain and spinal cord (TT1, Figure 2A, 2A’); jaw, branchial arches and pectoral fin buds (TT5, Figure 2B, 2B’); skeletal muscles (TT6, Figure 2C); central nervous system (TT7, Figure 2D); lateral line system (TT15, Figure 2E); rhombomere 5 (TT21, Figure 2F, 2F’); hindbrain and pectoral fin buds (TT28, Figure 2G, 2G’); retina (TT37, Figure 2H, 2H’); midbrain stripe (TT42, Figure 2I, 2I’) and spinal cord/pronephros (TT50, Figure 2J). For remaining lines, descriptions of their eGFP-rpl10a distribution patterns are shown in Supplementary Figures 1 and 2. All the information on the trap-TRAP transgenic lines has been uploaded to a web-based database: <u>trap-TRAP database</u>. This website displays TT lines pictures and descriptions of their expression pattern at different developmental stages, together with complementary information about the trap-TRAP technology.

**Figure 2.**
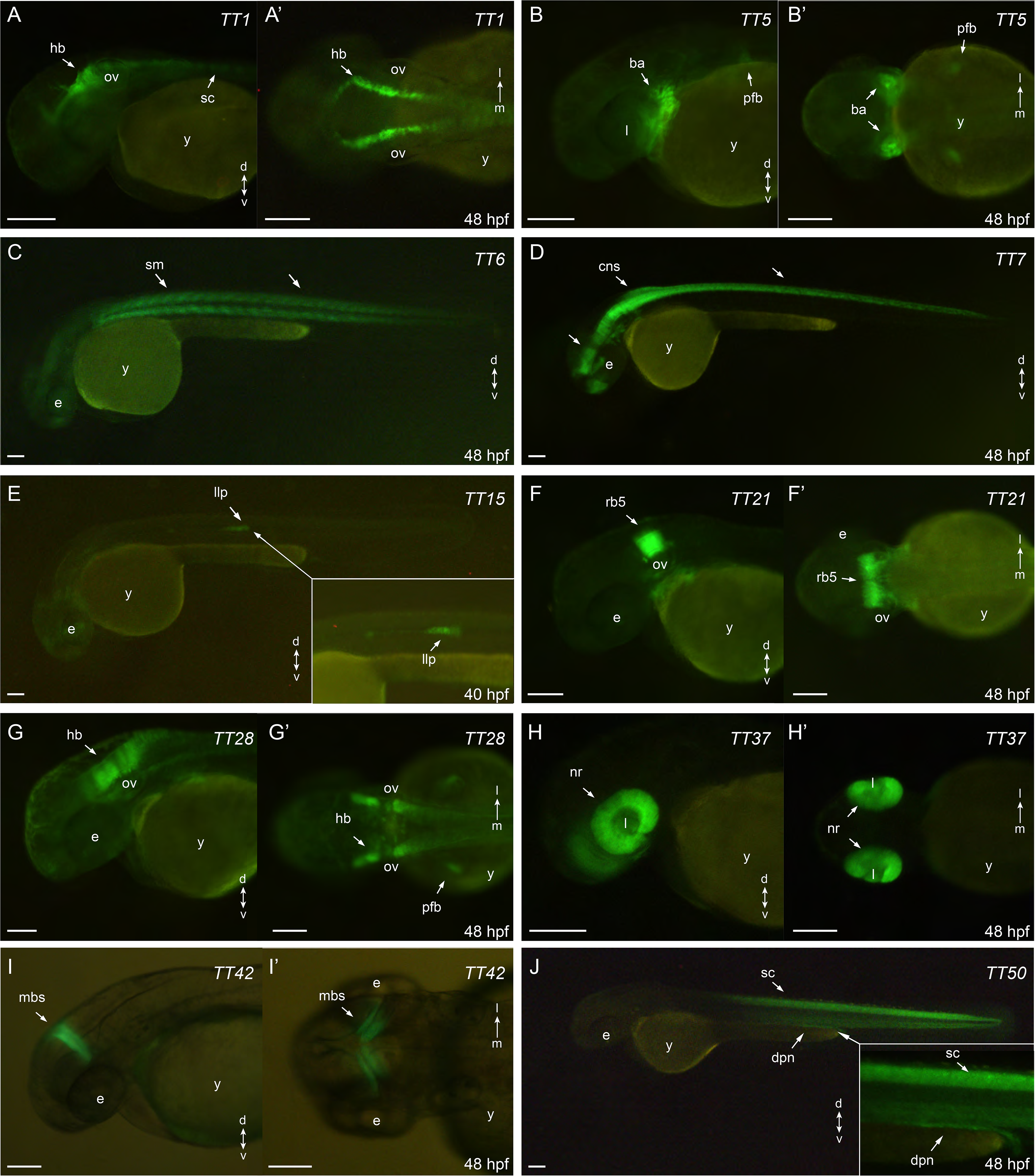
eGFP-rpl10a expression patterns: A, A’: TT1, hindbrain and spinal chord (lateral and dorsal, respectively); B, B’: TT5, jaw, brachial arches and pectoral fin buds (lateral and ventral, respectively); C: TT6, skeletal muscles (lateral); D: TT7, central nervous system (lateral); E: TT15, lateral line system (lateral); F, F’: TT21, rhombomere 5 (lateral and dorsal, respectively); G, G’: TT28, hindbrain and pectoral fin buds (lateral and dorsal, respectively); H, H’: TT37, retina (lateral and ventral, respectively); I, I’: TT42, midbrain stripe (lateral and dorsal, respectively); J: TT50, spinal cord and pronephros (lateral). ba = branchial arches; cns = central nervous system; dpn= distal pronephros; e = eye; hb =hindbrain; l = lens; llp =lateral line primordium; mbs= midbrain stripe; nr = neural retina; ov = otic vesicle; pfb = pectoral fin buds; rb5= rhombomere 5; sc = spinal cord, sm = skeletal muscles; y = yolk. Scale bar = 100 μm.

### Identification of trap-TRAP insertion sites

To investigate if the random insertions of the trap-TRAP cassette in the genome recapitulate the activity of nearby cis-regulatory elements, we mapped the targeted loci of a few representative lines by inverse PCR and compared their expression patterns with those of neighbouring genes. The analysis provided conclusive mapping results for 3 out of 4 lines examined (See Supplementary Table 1), the lines TT15, TT21 and TT42. In these cases, we successfully identified insertion sites in the vicinity of genes with an expression similar to that found for the corresponding strains (Figure 3). The line TT15, which showed expression in the lateral line, contains an insertion in the chromosome 1 (Figure 3A, 3A’). This region is 5’ distal to *lef1*, whose expression pattern includes a lateral line domain similar to that observed in TT15 embryos ^29^. Similarly, the analysis carried out on the TT21 line, which shows expression in rhombomere number 5, indicated that the insertion occurred on chromosome 23 (Figure 3B, 3B’). In the vicinity of the integration site is *mafba*, whose expression has been described in rhombomeres 5 and 6 ^30^. Finally, examination of the TT42 line identified an insertion on chromosome 8 (Figure 3C). This line shows a very specific expression stripe restricted to the midbrain (Figure 3C’). Accordingly, the neighbouring gene *her3* also shows expression in this domain ^31^. In all cases, the integration loci were also compared with data on chromatin accessibility, (ATAC-seq), as well as H3K27ac and H3K4me3 epigenetic marks, at 48hpf ^32, 33^. These comparisons suggest that the eGFP-rpl10a cassette could capture the activity of nearby cis-regulatory elements.

**Figure 3.**
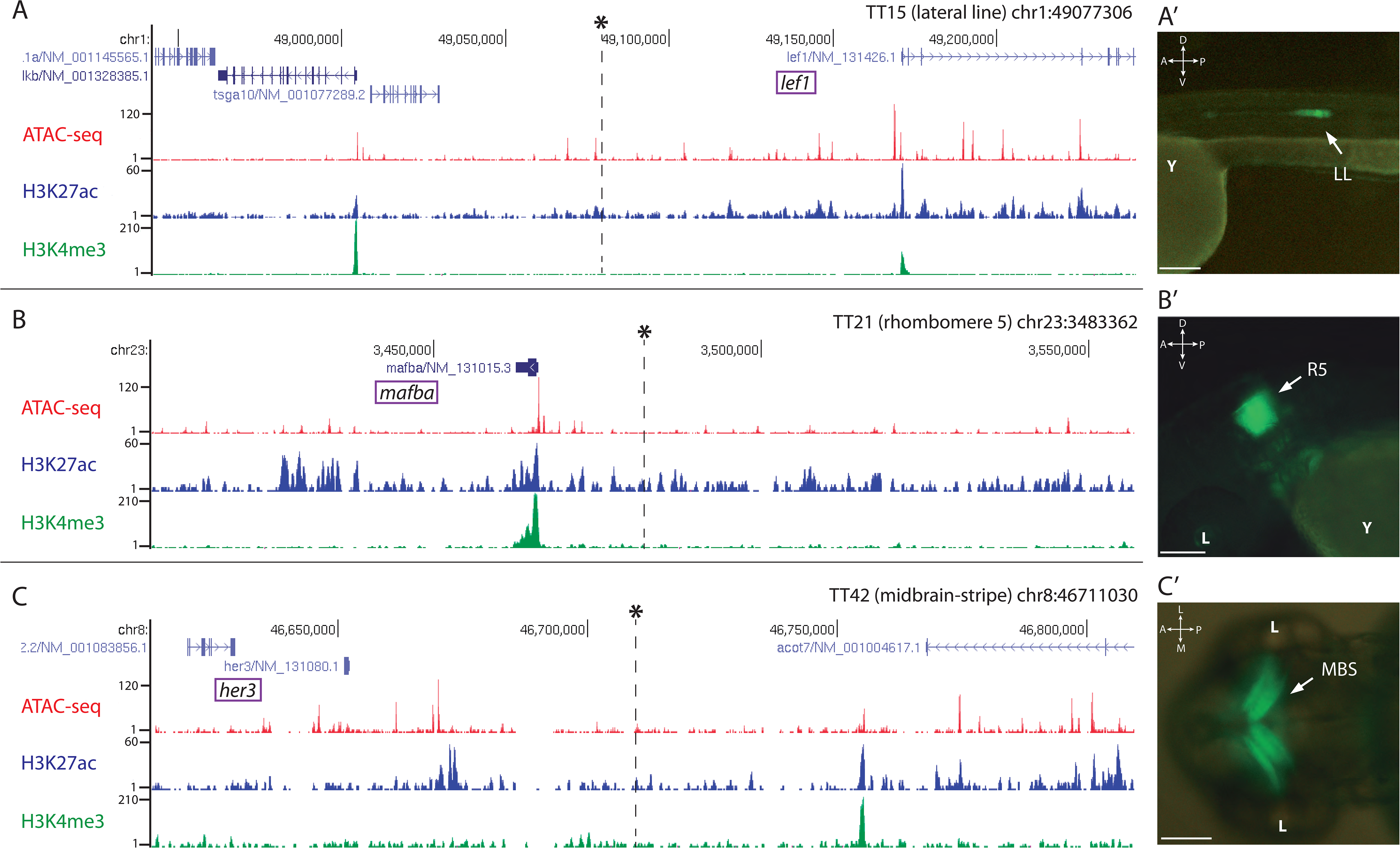
Genomic insertion sites of trap:TRAP cassette in representative transgenic lines. Insertions are marked with an asterisk*. Tracks for ATAC-seq, H3K27ac and H3K4me3 at 48 hpf are also shown for the different loci. A: Genomic location of the insert in the transgenic line TT15 in the vicinity of *lef1*, A’: TT15 expression pattern at 48 hpf. B: Genomic location of the insert in the transgenic line TT21 nearby *mafba*, B’: TT21 expression pattern at 48 hpf. C: Genomic location of the insert in the transgenic line TT42, C’: TT42 expression pattern at 48 hpf. LL= lateral line; R5= rhombomere 5; MBS= midbrain stripe. D-V= dorso-ventral; A-P= anterior-posterior. Scale bar = 100 μm

### Compatibility of TRAP technology with the Gal4/UAS system

As an alternative strategy to direct *eGFP*-*rpl10a* expression to tissues or cell types of interest, we combined the TRAP methodology with the Gal4-UAS system, which allows the spatiotemporal control of gene expression ^22, 34, 35^. To this end, the fusion gene was placed under the control of the UAS element to generate the *Tol2*_*UAS:TRAP* vector (Figure 1C). This construct was injected in one-cell stage zebrafish embryos together with *in vitro* -synthesized Tol2 transposase mRNA. To screen for potential adult founders, F0 animals were outcrossed with fish from a Gal4-expressing strain: *Tg[Rx3:Gal4]*, whose expression is restricted to the retina as previously reported ^36, 37^. As a reference control, we first verified retinal expression in the progeny of a *Tg[Rx3:Gal4]* × *Tg[UAS:RFP]* cross (Figure 4A). Then this retina-specific pattern was further confirmed in the founders progeny crossed to the Gal4 reference line: *Tg[Rx3:Gal4]* × *Tg[UAS:TRAP]* (Figure 4B). The fact that we did not observe toxicity, optic cup malformations, or developmental delays in these crosses indicates that Gal4 drivers can be successfully combined with the UAS:TRAP line to specifically perform TRAP analysis in a broad variety of tissues and cell types in zebrafish.

**Figure 4.**
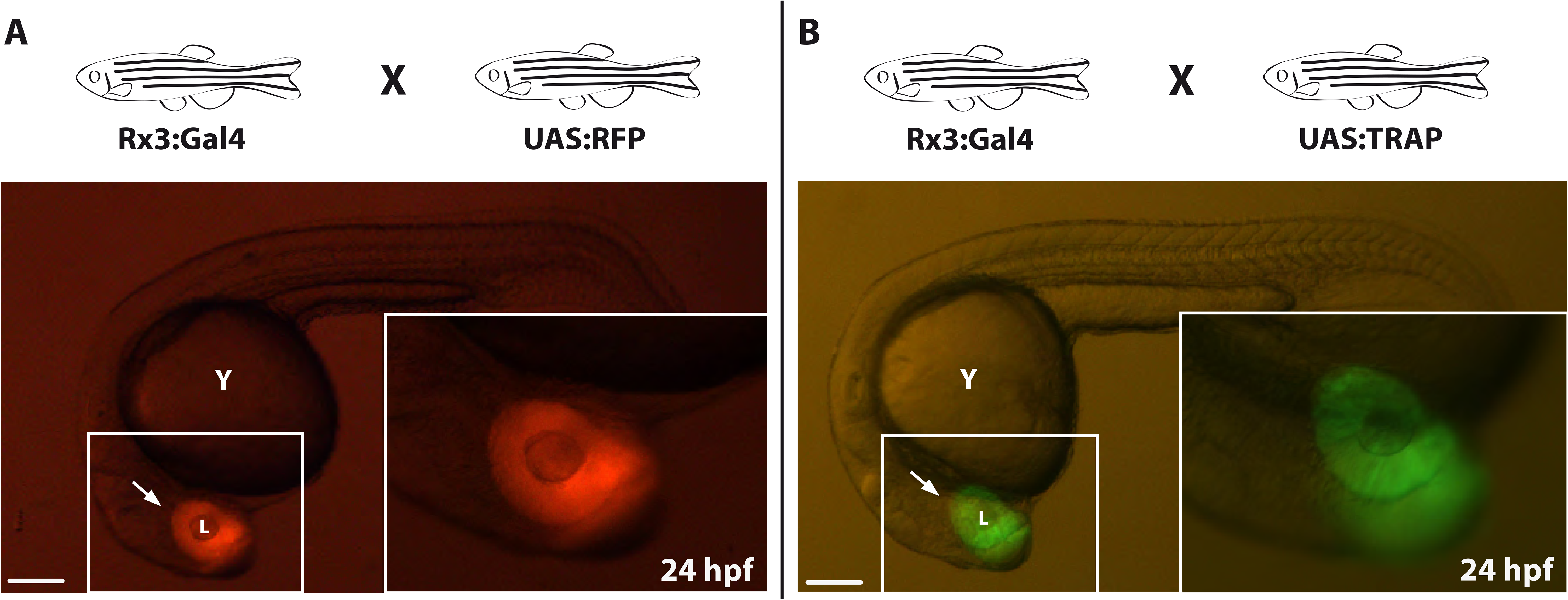
Compatibility of TRAP technology with the Gal4/UAS system. A: Zebrafish embryo from the control cross *Tg[Rx3:Gal4]* × *Tg[UAS:RFP]* showing RFP expression in the developing retina (arrow) at 24 hpf. B: zebrafish embryo derived from a *Tg[Rx3:Gal4]* × *Tg[UAS:TRAP]* cross showing *eGFP*-*rpl10a* expression at 24 hpf in the retina (arrow). l = lens; y = yolk. Scale bar = 100 μm.

## Conclusions

We have described two different approaches, Trap-TRAP and UAS-TRAP, which allow expanding the use of the TRAP methodology in zebrafish and can be easily adapted to other model organisms. While TRAP methods allow tissue-specific isolation of the mRNA fraction being translated, a closer representation of the protein profile of the cell at a given time ^13^, the need to generate transgenic lines for each type of analysis has limited their application. Our strategies facilitate the fast and efficient generation of transgenic zebrafish strains suitable for TRAP analysis. Previous studies in zebrafish have reported the use of TRAP methods for translational profiling in zebrafish using a few tissue-specific promoters ^14, 15, 16, 17^. Here we applied the Trap-TRAP strategy to generate random insertions of the *eGFP*-*rpl10a* cassette into the genome. In our pilot screen we examined 53 founders, 33 of them giving rise to tissue-specific stables lines, in whose progeny no developmental defects were observed, highlighting the compatibility of our approach with the large-scale generation of TRAP lines. Using a strategy similar to that of a previous study in *Drosophila* ^12^, we also generated and tested a UAS:TRAP transgenic line that will allow to combine the TRAP methodology with the large collection of Gal4 lines currently existing in zebrafish. These two alternative approaches can facilitate the systematic generation of transgenic lines available for TRAP analysis. A potential limitation of them could be the sensitivity of the TRAP method when small cell populations are interrogated or weak promoters are employed. However, the access to large numbers of synchronized embryos (i.e. once the stable lines have been generated), together with evidence from previous TRAP analyses in zebrafish targeting populations limited in number ^17^, suggest that in many cases this should not be a major obstacle. A further refinement of the Trap-TRAP and UAS-Trap approaches may include the generation of transgenic lines harbouring an Avi-tagged *eGFP*-*rpl10a* cassette, as previously described ^16^. This system, in combination with BirA activating strains, does not depend on the use of specific antibodies for immunoprecipitation of tagged ribosomes and may result in an increased sensitivity.

## Materials and Methods

### Fish maintenance

The zebrafish (*Danio rerio*) AB/Tübingen (AB/TU) wild-type strain, and the transgenic line *Tg[Rx3:Gal4:UAS:RFP]* ^36^ were maintained under standard conditions at 28°C. Embryos and larvae were kept in E3 medium (5 mM NaCl, 0.17 mM KCl, 0.33 mM CaCl2, 0.33 mM MgSO4) supplemented with Methylene Blue (Sigma) at 28°C and staged according to somite number and morphology ^38^. Animal experiments were carried out according to ethical regulations. Experimental protocols have been approved by the Animal Experimentation Ethics Committees at the Pablo de Olavide University and CSIC (license number 02/04/2018/041).

### TRAP vectors construction

To develop the different approaches carried out in this work, two different constructs were built, based on the *Tol2* transposition system ^28^: *Tol2*_*trap:TRAP* and *Tol2*_*UAS:TRAP*. Both constructs were obtained through modifications of the Tol2-zTRAP plasmid, which contains the eGFP-rpl10a fusion gene flanked by Tol2 sites ^14^. The *Tol2*_*trap:TRAP* construct was generated by inserting the *gata2p* minimal promoter from the ZED vector ^25^ into the *Tol2*-*zTRAP* multiple cloning site using *SalI* and *BamHI* restriction enzymes. The *Tol2*_*UAS:TRAP* plasmid was built by inserting the UAS fragment from the Tol2kit p5E_UAS plasmid (Kwan et al., 2007) into the *Tol2*-*zTRAP* multiple cloning site using *BamHI* and *NheI*.

### Transgenic lines generation

The different transgenic lines were generated on the (AB/TU) background by microinjection of each vector (*Tol2*_*trap:TRAP* or *Tol2*_*UAS:TRAP*) into one-cell stage embryos. Following the Tol2 transposon/transposase transgenesis method ^20^, 100-200 pg of the plasmid were injected together with 100-200pg of the *Tol2* transposase mRNA.

#### Trap-TRAP lines

Embryos microinjected with the *Tol2*_*Trap:TRAP* construct were examined at 24 hpf and those showing any type of fluorescence were raised. When the selected fish reached adulthood, they were outcrossed with wild-type animals, and their progeny was examined at 24, 48 and 72 hours post-fertilization (hpf) in order to identify and characterize the different *eGFP:rpl10a* expression patterns. Positive transgenic fish were considered as founders, and their progenies were raised to expand the different lines. In some cases, embryos from the same parental fish display different expression patterns, as a result of multiple integrations. In these cases, the different expression patterns were isolated through outcrossing and each different embryo was raised as an independent line. The integrity of the *eGFP:rpl10a* cassette at the landing loci was assessed by PCR using the specific primers TrapS_fwd: 5’-CATGTCGACAAGTGTCCG, and TrapS_rev: 5’-TGCATTCTAGTTGTGGTTTGTCC).

#### UAS:TRAP lines

Embryos microinjected with the *Tol2*_*UAS:TRAP* construct were also screened once they reached adulthood. To test the construct insertion and its functionality, microinjected fish were crossed with *Tg[Rx3:Gal4]* fish. To assess the expression pattern resulting from this cross, *Tg[Rx3:Gal4;UAS:TRAP]* embryos were compared with reference embryos from the *Tg[Rx3:Gal4;UAS:RFP]* line ^36^.

### Imaging

The *eGFP:rpl10a* expression pattern displayed by the embryos of each trap-TRAP line was examined and photographed using an Olympus fluorescence micro stereoscope equipped with a camera Nikon (). Pictures were taken at 24, 40, 48 and 72 hpf. To prevent embryo pigmentation and facilitate expression pattern analysis, E3 medium was supplemented with 0.2 mM 1-phenyl-2-thiourea (PTU). In the case of the *UAS:TRAP* and the different Gal4 strains crosses, pictures of the progeny were taken at 24 hours post fertilization. In all cases, the embryos were dechorionated with forceps if needed and mounted in 2% methylcellulose.

### Identification of Tol2_Trap:TRAP integration sites

From each analysed line (TT15, TT21, TT37 and TT42), genomic DNA was isolated from 5 individual 72hpf zebrafish embryos using Chelex 100 sodium form (Sigma Aldrich). Briefly, embryos were incubated in 45μL of Chelex 5% together with 5 μL of proteinase K (10mg/mL) for a minimum of 2 hours. Afterwards, proteinase K was heat-inactivated (95°C, 10 min.). To prepare PCR templates, 5 μL (approximately 250 ng) of the extracted DNA was first digested using the restriction enzyme DpnII (NEB), diluted to 100 μL and ligated with T4 ligase (NEB). The resultant ligation was used as a template for inverse PCRs. Two different nested PCRs were performed to amplify the junction fragments containing both Tol2 ends and their adjacent genomic DNA. The first PCR product was diluted to 200 μL with ddH2O before the second PCR. The primers and PCR programs used were described in a previous study ^39^. The common band amplified from the five embryos was purified and cloned in TOPO (Invitrogen, pCR8/GW/TOPO TA cloning KIT). The resulting clones were sequenced using the T7 primer and the sequences obtained were subsequently examined to identify the adjacent fragments, which were aligned to the zebrafish genome (GRCz10/danRer10) using the UCSC BLAT Search Genome tool (https://genome.ucsc.edu/cgi-bin/hgBlat). The expression profile of nearby genes was then compared them with that of the corresponding line.

### Website

All information on the different trap:TRAP lines was uploaded to the zebrafish TRAP database website: <u>trap-TRAP database</u>: (https://amfermin.wixsite.com/website). Photographs of the different trap:TRAP lines at different developmental stages were included together with their descriptions and the trap:TRAP methodology.

## Acknowledgements

We thank Rocío Polvillo for their excellent technical assistance. This work is supported by grants awarded to JRMM from the Fundación Ramón Areces (program-2016); PY20_00006 from Junta de Andalucía; as well as Spanish Ministry of Science, Innovation and Universities (MICINN, AEI/FEDER) BFU2017-91324-EXP, BFU2017-86339P, RED2018-102553-T, PID2020-112566GB-I00 and MDM-2016-0687.

## Author contribution

JC conducted most experiments and analyses with the help of ESR. AFM contributed to the design and construction of the webpage. JRMM conceived the project and assisted JC in data analysis. The manuscript was edited and written by JC and JRMM.

## Conflict of Interest

The authors declare that they have no conflict of interest

## Supplementary Figure Legends

**Figure Supplementary 1.**
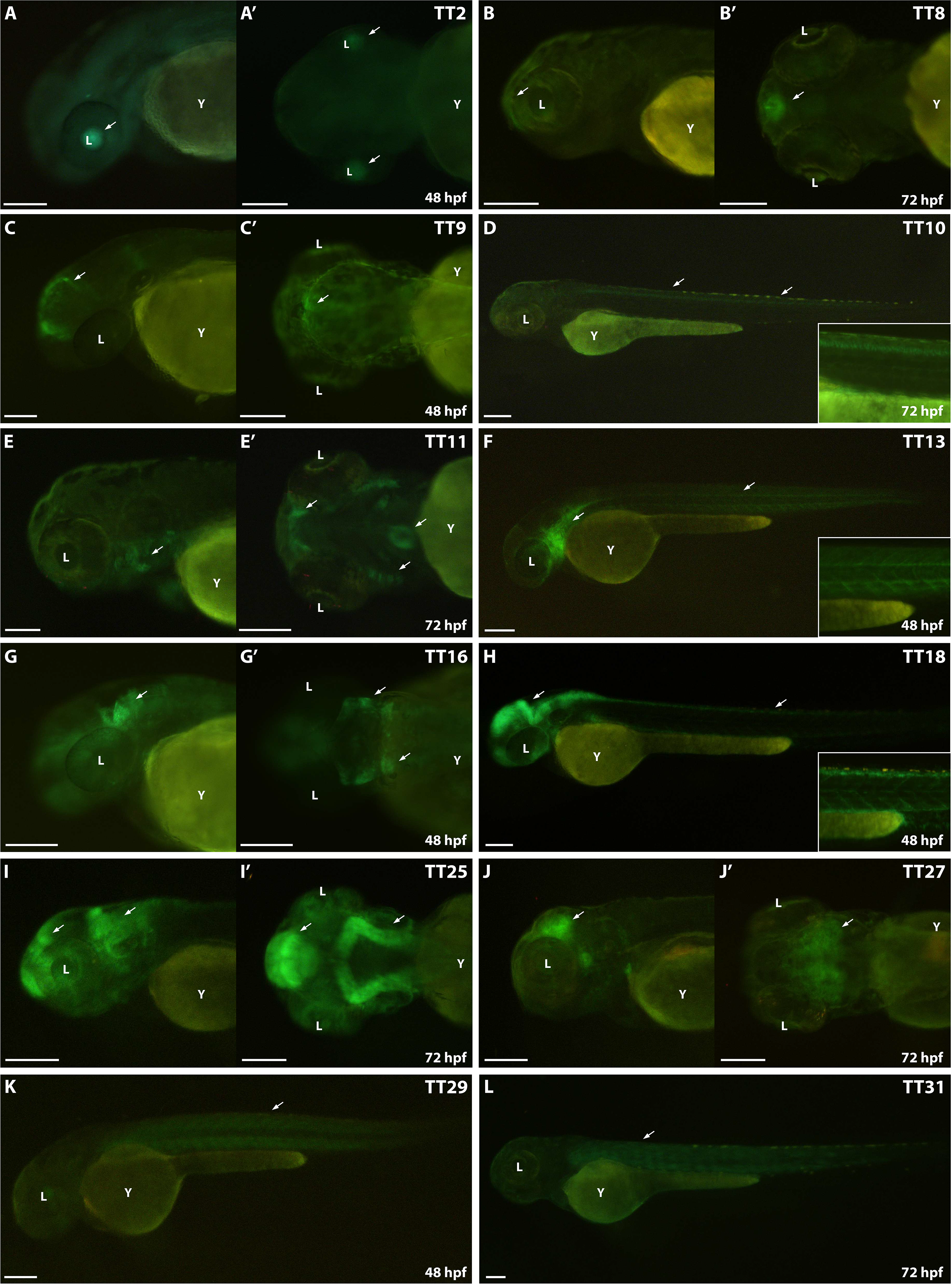
eGFP-rpl10a expression patterns (TT2 - TT31). **A, A’**: TT2, lenses (lateral and dorsal, respectively); **B, B’**: TT8, forebrain, eyes (lateral and ventral, respectively); **C**, **C’**: TT9, dorsal optic tectum (lateral and dorsal, respectively); **D**: TT10, spinal chord neurons (lateral); **E**: TT11, branchial arches, jaw and heart (lateral and ventral respectively); **F, F’**: TT13, vascular system (lateral); **G, G’**: TT16, anterior hindbrain (lateral and dorsal, respectively); **H, H’**: TT18, Central nervous system, pectoral fin buds (lateral); **I, I’**: TT25, dorsal central nervous system, eye (lateral and dorsal, respectively); **J, J’**: TT27, midbrain (lateral and dorsal, respectively); **K**: TT29, lenses, skeletal muscles and central nervous system (dorsal); **L**: TT31 skeletal muscles (dorsal). l= lens; y = yolk. Scale bar = 100 μm.

**Figure Supplementary 2.**
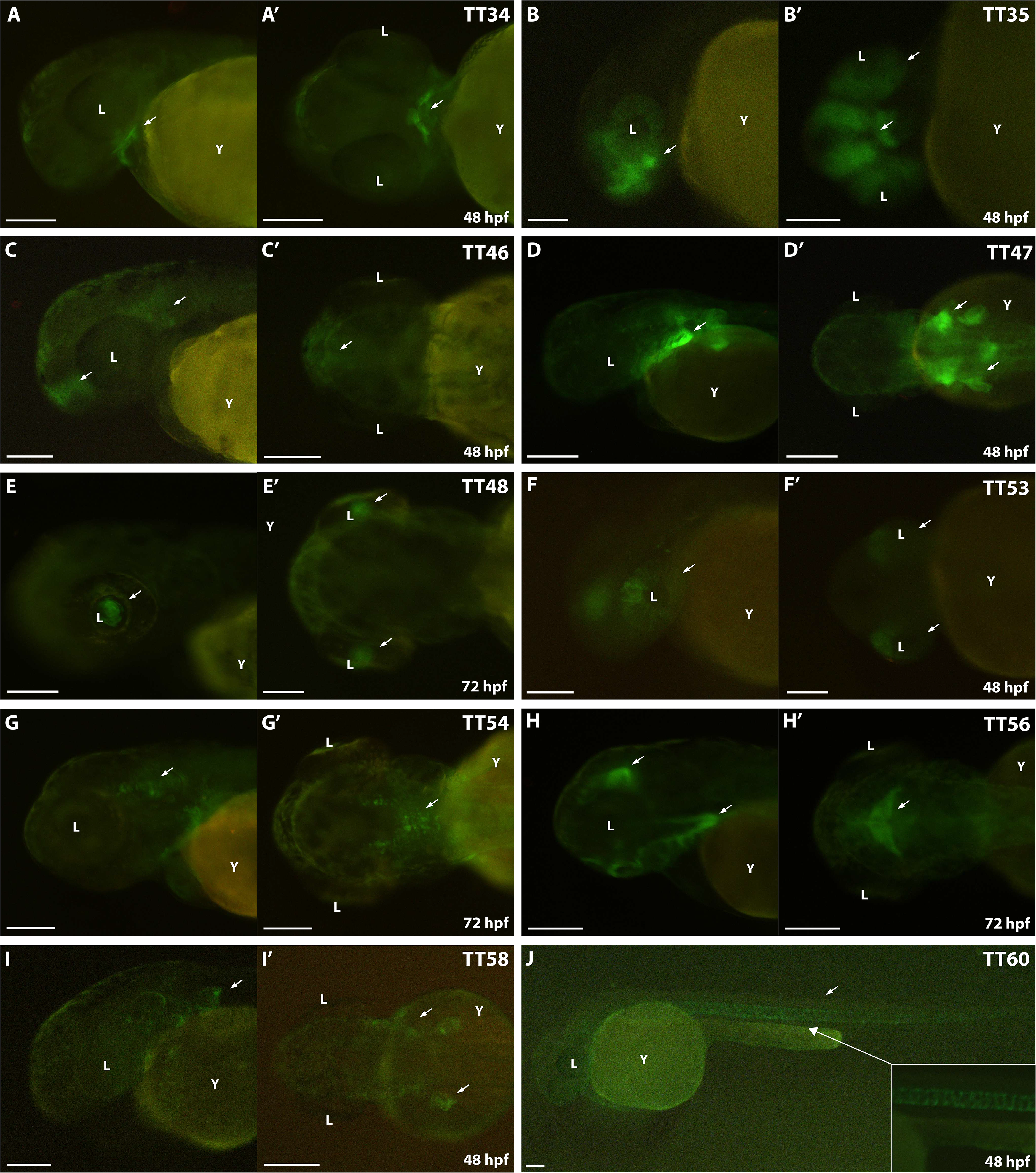
eGFP-rpl10a expression patterns (TT34 - TT60). **A, A’**: TT34, heart (lateral and dorsal, respectively); **B, B’**: TT35, telencephalon, eyes (lateral and ventral, respectively); **C, C’**: TT46, telencephalon (lateral and dorsal, respectively); **D, D’**: TT47, pectoral fin buds, branchial arches (lateral and dorsal, respectively); **E, E’**: TT48, lens fibers (lateral and ventral respectively); **F, F’**: TT53, dorsal retina (lateral and ventral, respectively); **G, G’**: TT54, ventral hindbrain (scattered cells), branquial arches (lateral and dorsal, respectively); **H, H’**: TT56, midbrain subdomain, branchial arches, jaw (lateral and dorsal, respectively); **I, I’**: TT58, pectoral fin buds, (lateral and dorsal, respectively); **J**: TT60, notochord (lateral); l= lens; y = yolk. Scale bar = 100 μm.

**Figure Supplementary 3.**
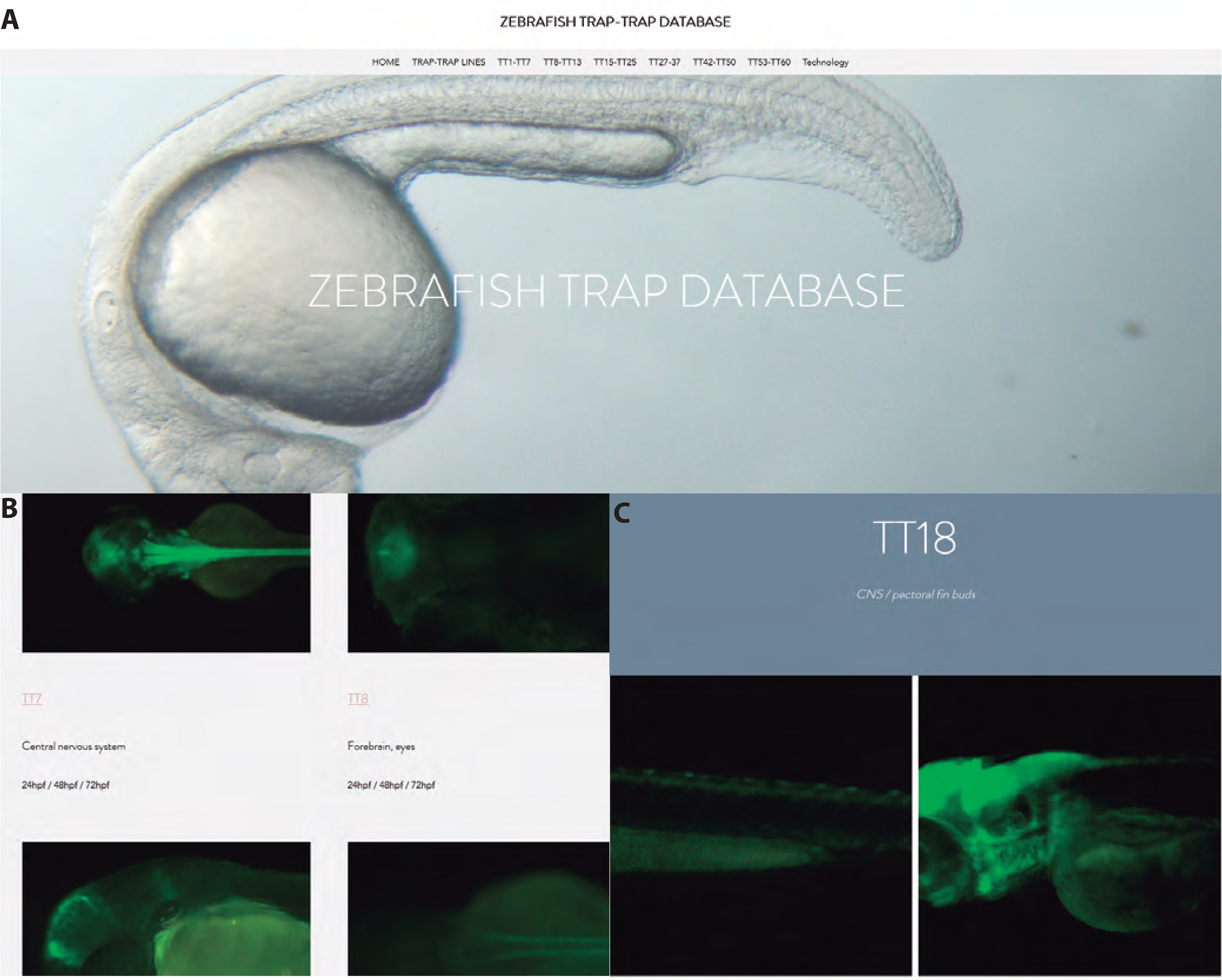
Content of the zebrafish trap-TRAP database. (A) Homepage of the website. The menu at the top contains direct links to the different pages of the site. (B) Images of the summarizing page “trap-TRAP lines” with descriptions and links to each of the trap-TRAP lines. (C) Specific example for the tab displaying the TT18 line. Each tab shows a galary of images illustrating the expression pattern of an individual line.

**Table S1.**
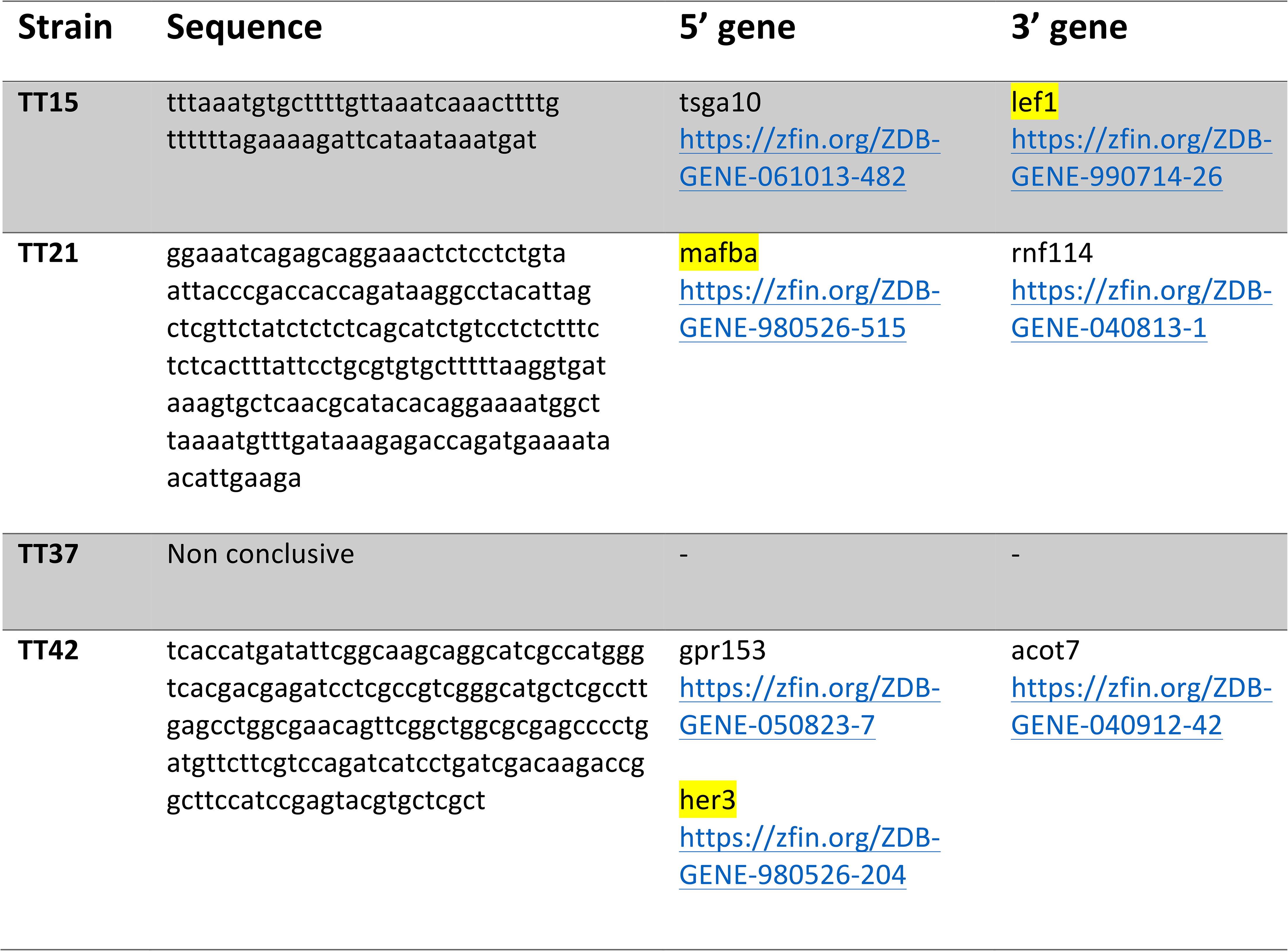
Analysed strains, corresponding cloned sequences matching genomic positions, and nearby 5’and 3’ genes.

## Notes

### Competing Interest Statement

The authors have declared no competing interest.

